# Transformer Language Models Reveal Distinct Patterns in Aphasia Subtypes and Recovery Trajectories

**DOI:** 10.64898/2026.03.27.714240

**Authors:** Seyed Saeed Ahmadi, Julius Fridriksson, Dirk B. den Ouden

**Author notes:** Corresponding author. S. Saeed Ahmadi, Department of Communication Sciences and Disorders, University of South Carolina, Columbia, SC, USA.

## Abstract

Language impairments in aphasia are characterized by various representational disruptions that may be reflected in discourse production. This research examines the capacity of transformer-based language models, particularly GPT-2, to serve as a computational framework for analyzing variations in aphasic narrative speech. A longitudinal dataset of narrative speech samples collected at six time points from individuals with aphasia (N = 47) was utilized as part of an intervention study. All transcripts were processed via the GPT-2 language model to obtain activation values from each of the 12 transformer layers. Statistically significant differences in activation magnitude across aphasia subtypes were found at every layer (all p < .001), with the most pronounced effects in the deeper layers. Pairwise Tukey HSD tests revealed consistent distinctions between Broca’s aphasia and both Anomic and Wernicke’s aphasia, suggesting a shared activation profile between the latter two. Longitudinal tests revealed significant changes over time, especially in the final three layers (10–12). These findings suggest that transformer-based activation patterns reflect meaningful variation in aphasic discourse and could complement current diagnostic tools. Overall, GPT-2 provides a scalable tool to model representational dynamics in aphasia and enhance the clinical interpretability of deep language models.

## Introduction

Aphasia is an acquired language disorder that typically occurs after a stroke and significantly affects communication, quality of life, and social participation (Brady et al., 2016). While lesion characteristics, neuronal integrity, and treatment modalities affect the intricate process of aphasia recovery, scalable and reliable methods for monitoring language recovery remain limited.

Aphasia is not a homogeneous condition; it comprises various subtypes that differ in clinical presentation and neural underpinnings (Zhao et al., 2023). The classification of aphasia types remains controversial. Broca’s aphasia is a nonfluent type of aphasia characterized by agrammatic speech production, although comprehension remains generally intact (Hickok, 2009). Wernicke’s aphasia is characterized by fluent yet semantically empty speech and compromised comprehension (Potagas, 2011). Other subtypes of aphasia further highlight the heterogeneity of language deficits following stroke. It is important to consider these distinctions, as collapsing across subtypes may obscure clinically meaningful variation in recovery patterns and neural reorganization. By explicitly examining differences within subtypes, studies can identify whether computational markers generalize across aphasia or reflect subtype-specific impairment, ultimately supporting more accurate diagnostics and individualized treatment planning.

Recent advancements in artificial intelligence, particularly in natural language processing (NLP), have introduced novel methods for characterizing language deficits. NLP enables computational analysis of fluency, coherence, and syntax in clinical speech, offering new avenues for assessing language impairments, and progress in this area has accelerated with the development of large language models. Unexpectedly, the internal representations of these models have even been found to align with human brain activation during language processing (Caucheteux et al., 2022; Caucheteux & King, 2022; Gleichgerrcht et al., 2021; Schrimpf et al., 2021).

Transformers, especially models such as GPT-2, process language via multi head self-attention and deep hierarchical structures that encode linguistic and semantic information across layers (Vaswani et al., 2017; Yeh et al., 2023). Different attention heads and layers have been suggested to specialize in particular linguistic phenomena (Vig & Belinkov, 2019). The architectural characteristics of the GPT-2 make it a valuable instrument for investigating language impairments in aphasia at a representational level, given the demonstrated correlations between layer activation in the GPT-2 and neural activity (Caucheteux et al., 2022). In this context, the output value that each layer produces in response to linguistic input is referred to as its “activation,” which reflects the magnitude of that layer’s response and the extent to which it encodes aspects of the text’s structure or meaning. Previous research has demonstrated that deeper layers reflect higher-level semantic and discourse-level information, whereas surface layers are more sensitive to lexical and syntactic characteristics (Geva et al., 2020; Jawahar et al., 2019; Tenney et al., 2019). Importantly, human programmers have not specifically coded these levels to encode certain linguistic functions. During training on large-scale text corpora, the model adjusts its internal weights to capture statistical regularities of language. As a result, any specialization observed in a particular layer is not predefined but emerges through learning (Geva et al., 2020; Rogers et al., 2021).

Within this architecture, large language models encode text across successive levels, where representations in different layers may more strongly reflect distinct linguistic features such as lexical, syntactic, or semantic information. The surface layers primarily address superficial patterns, including word forms, punctuation, and basic part-of-speech tags. The middle layers begin to encode grammatical structures, such as rules, clause demarcations, and sentence constructions, such as how one might evaluate phrase structure in continuous speech. The deeper layers emphasize semantic significance and discourse-level characteristics, addressing broader context, reference resolution, and speaker purpose (Jawahar et al., 2019; Peters et al., 2018). This tiered organization provides a framework for interpreting layer wise analyses of aphasic and typical speech.

The OpenAI GPT structure is based on multi head self-attention (Radford et al., 2018). A key advantage of self-attention is its interpretability, which allows researchers to examine which input elements are treated as most relevant for text parsing (Belinkov & Glass, 2019). The small version of GPT-2 includes 12 layers and 12 attention heads.

In this study, we aimed to investigate whether the internal layer activation patterns of GPT-2 differ between individuals with various types of aphasia and neurotypical individuals. Specifically, we examine (1) whether the representational patterns in aphasic speech differ from typical (unimpaired) language patterns, (2) whether these differences vary across aphasia subtypes, and (3) how these patterns evolve during language recovery.

If significant associations are discovered between large-language-model activation patterns and standardized language assessment scores, these patterns could be used as computational biomarkers for aphasia. Recent work has shown that surprise-based indices from GPT-style models discriminate aphasia presence and subtype more accurately than traditional linguistic measures do (Cong et al., 2024). By obtaining these markers from precise, naturalistic speech samples, clinicians could (i) create objective, low-effort tools for diagnosis and monitoring, (ii) enhance subtype classification for personalized treatment, and (iii) improve the prediction of therapy responsiveness. Furthermore, by bridging the gap between neurolinguistics and language model studies, this research contributes to the theoretical understanding of the reconfiguration of language following a stroke.

We analyzed narrative speech collected from an intervention study in individuals with aphasia over multiple time points, extracting GPT-2 hidden-state activations. Statistical and machine learning techniques were used to evaluate longitudinal changes and classify aphasia subtypes on the basis of model-derived representations.

## Methods

This study used the GPT-2 across six time points to examine longitudinal changes in internal language representations in individuals with aphasia enrolled in an intervention study aimed at identifying predictors of response to treatment (POLAR dataset (Kristinsson et al., 2023)).

### Participants

The participants were selected from a larger dataset, which included 102 individuals with poststroke aphasia (Kristinsson et al., 2023). From this group, we included the 57 individuals who produced at least 100 words in their speech sample at the first time point. In addition, we included 10 non-aphasic participants without a history of stroke who served as neurologically typical controls. For the longitudinal analyses, a subset of 47 aphasic individuals was selected on the basis of having sufficient narrative output (at least 100 words per story) and attending assessment sessions at all six time points. The participant demographics and clinical characteristics are presented in Appendix A, Table A1.

### Narrative Elicitation and Transcription

Narrative discourse samples were collected at six time points: (1) baseline (pretreatment), (2) following the initial treatment phase, (3) post rest period, (4) after the second treatment phase, (5) one-month post-treatment, and (6) six months post-treatment. At each time point, participants were asked to retell the story of *Cinderella* using a wordless picture book to ensure standardized elicitation (Saffran et al., 1989). All audio samples were transcribed manually by trained graduate students in speech-language pathology following the AphasiaBank transcription protocol (Macwhinney et al., 2011), although only textual transcription (without error coding) was used in this study. Transcripts were reviewed for accuracy and consistency, and all words were converted to lowercase to provide uniformity in input to the model. The finalized transcripts were tokenized via the GPT-2 tokenizer.

### Language Model and Feature Extraction

We employed the pretrained GPT-2 small model (12 layers) to analyze the transcribed narratives. For each transcript, we extracted hidden state activations from all 12 layers of GPT-2 for every token, controlling the number of tokens so that all transcripts had the same number of tokens. Token-level activations were averaged within each layer to yield a 768-dimensional vector per layer per sample, representing the internal linguistic representation of each narrative at each time point.

### Data analysis

We implemented a multitiered statistical framework via R (Team, 2025). Our analysis pipeline consisted of six steps:

#### 1. Baseline Group Comparison of GPT-2 Layer Activations

To investigate whether baseline layer wise hidden-state activations of the GPT-2 differ between individuals with aphasia and neurotypical controls, we extracted mean activation values from each of the 12 transformer layers for all participants for all Cinderella narrative discourse samples. We conducted nonparametric Wilcoxon rank-sum tests for each layer to compare the activation distributions across the two groups. This method enabled us to ascertain whether layers in the GPT-2 model encapsulated group-specific linguistic representations at the baseline point.

#### 2. Layer Wise Activation Differences across Aphasia Subtypes at Baseline

To examine whether various subtypes of aphasia displayed unique representational profiles at baseline in the GPT-2, we performed a layer wise analysis of hidden-state activations at baseline (Time Point 1). Kruskal–Wallis rank-sum tests were conducted for each of the 12 transformer layers to assess variations in the mean activation values across five aphasia subtypes: Broca’s, Wernicke’s, Conduction, Anomic, and Global aphasia, as defined with the WAB-R (Kertesz, 2007), and one control group. Dunn’s post hoc paired tests, utilizing the Benjamini–Hochberg correction, were subsequently employed for layers exhibiting significant group effects to ascertain which subtype pairs contributed to the observed variation.

#### 3. Group-level activation profile

To examine representational patterns in aphasic speech, we conducted a group-level analysis of GPT-2 hidden-state activations. At each time point (1–6) and for each of the 12 model layers, we extracted token-level hidden states from the transcripts of 47 individuals with aphasia. These activations were first averaged within each transcript to obtain a transcript-level mean. Next, transcript-level means were averaged across participants, resulting in a 6 (time points) × 12 (layers) matrix of mean activation values. In addition, to statistically assess changes in activation over time and across model layers, we fitted a linear mixed-effects model with participant-level random intercepts:

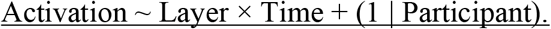

#### 4. Layer Activation Patterns across Aphasia Subtypes

To examine variation in representational patterns across standard aphasia subtypes, we computed mean GPT-2 hidden state activations separately for each subtype group, as defined with the WAB-R. To statistically evaluate group-level differences across aphasia subtypes, we conducted twelve one-way ANOVAs (one per layer), with the aphasia subtype as the between-subjects factor and the mean activation value at each layer at each time point as the dependent variable. Post hoc comparisons were performed via Tukey’s HSD test.

#### 5. Longitudinal Dynamics of GPT-2 Layer Activations over Time

To examine how representational patterns changed over time in response to aphasia therapy, we fitted linear mixed-effects models for each GPT-2 layer, using activation as the dependent variable and time point (1–6) as an ordinal independent variable. Aphasia subtypes were included as between-subject predictors, allowing us to test for interactions between subtypes and improvement trajectories. A random intercept for participants was included to account for within-subject repeated measures. This approach enabled us to assess longitudinal trends in activation while reducing the multiple comparison burden inherent in individual-level regressions. In addition, we used Wilcoxon signed-rank tests to compare time point 1 and time point 6 activation values separately for each of the 12 GPT-2 layers, assessing changes in layer–layer activation after treatment. This nonparametric test was chosen because of nonnormal distributions and the presence of outliers in the GPT-2 activation data. Given the use of only two time points and the absence of assumptions regarding linearity or equal variances, the Wilcoxon tests provided a more robust alternative to regression. Effect sizes (r) were computed for each layer-specific comparison, and the results were subsequently corrected for multiple comparisons to ensure statistical validity.

#### 6. Correlation with Clinical Language Scores

We assessed the relationship between GPT-2-layer activations and aphasia severity by calculating the average activation for each patient across all 12 transformer layers and correlating these values with their respective WAB-AQ scores. Layer wise correlation analyses were performed using the full range of WAB-AQ values to examine how activation patterns varied with aphasia severity.

To control for alpha inflation resulting from the large number of layer wise and group-level statistical tests conducted across analyses, all reported p values were adjusted via Benjamini– Hochberg false discovery rate (FDR) correction (Benjamini & Hochberg, 1995). Specifically, FDR adjustment was applied across the 12 GPT-2 layers within each analysis. This approach ensures that significant effects reflect robust differences rather than chance findings due to multiple comparisons.

## Results

### Baseline Group Comparison of GPT-2 Layer Activations

Wilcoxon rank-sum comparisons of GPT-2 hidden-state activations between the aphasia and control groups at Time Point 1 indicated that although the overall activation patterns were similar (with deeper layers showing greater activation than surface layers), there were statistically significant changes across layers 1, 2, 5, 8, 10, 11, and 12 of the transformer. Specifically, activations were higher in layers 12, 11, and 8 in the aphasia group than in the control group, whereas they were lower in layers 10, 5, 2, and 1 (all adjusted p-values < 0.05). This suggested that, at baseline, layer wise representational profiles for the Cinderella narrative differed between aphasic neurotypical participants.

### Layer Wise Activation Differences across Aphasia Subtypes at Baseline

Statistically significant variations in GPT-2 activations were detected among all groups throughout all 12 layers (adjusted p < .05), with notably pronounced effects in layers 5, 9, 10, 11, and 12. Post hoc analysis indicated that Broca’s aphasia consistently differed from the Anomic, Conduction, and Global subtypes across multiple deeper layers. Furthermore, differences between Anomic and Conduction aphasia were noted in layers 1, 5, and 9. Compared with non-aphasic subjects, only the Broca, Anomic, and Conduction aphasia subtypes presented considerable abnormalities, particularly in layers 5, 9, 10, and 11.

### Group-level activation profile

Averaged GPT-2 hidden-state activations were calculated across all patients for each model layer (1–12) and each time point (time points 1–6). These values are visualized in Figure 2 as a heatmap. Higher activation values were observed in deep layers across all time points. The overall layer wise activation profile remained consistent throughout the six-month period. To statistically quantify these observations, we fitted a linear mixed-effects model with a Layer × Time interaction. The analysis revealed a significant main effect of Layer (*F*(11, 3266) = 4847.8, *p* < 2×10^−16^), indicating substantial differences in activation magnitudes across the 12 layers. There was also a significant main effect of time (*F*(5, 3266) = 33.8, *p* < 2×10^−16^) and a significant layer × time interaction (*F*(55, 3266) = 18.9, *p* < 2×10^−16^), suggesting that temporal activation trajectories differed by layer. Post hoc analyses revealed that Layers 10, 11 and 12 had consistently greater activation and the greatest variability across time (Layer 12 = 0.0066, Layer 11 = 0.0049, Layer 10 = 0.0027).

**Figure 1.**
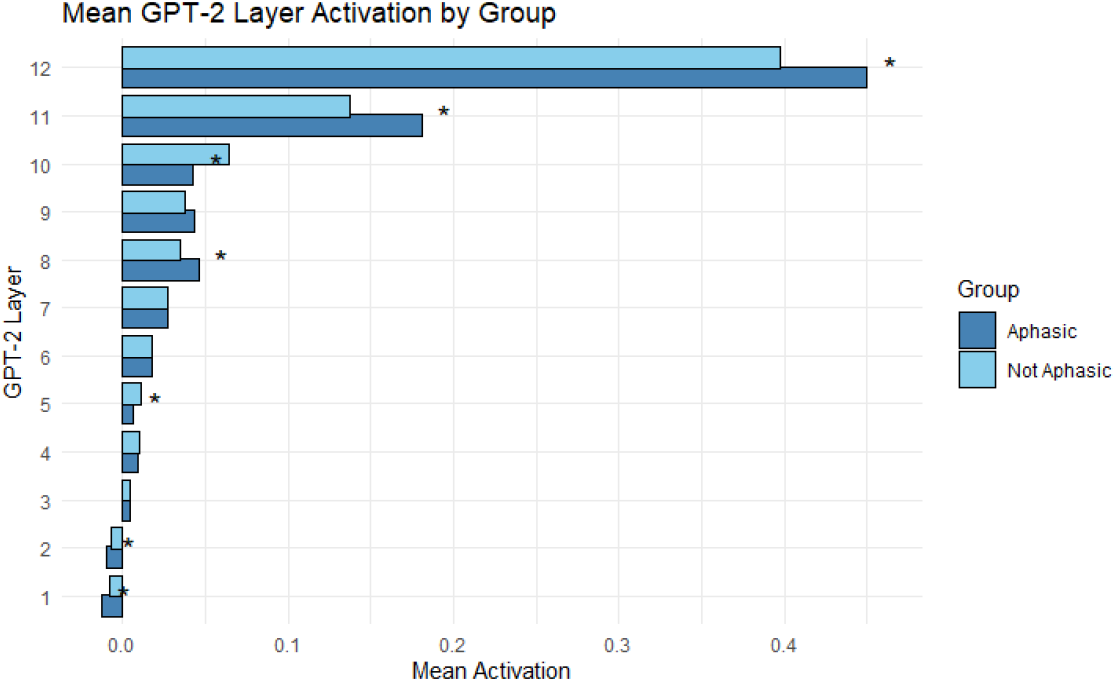
Mean GPT-2 Layer Activation in Aphasic and Non-Aphasic Groups. Bar plot showing the average GPT-2-layer activation across 12 layers for individuals with aphasia (blue) and healthy controls (skyblue). Each bar represents the mean activation value for one group at each layer.

**Figure 2.**
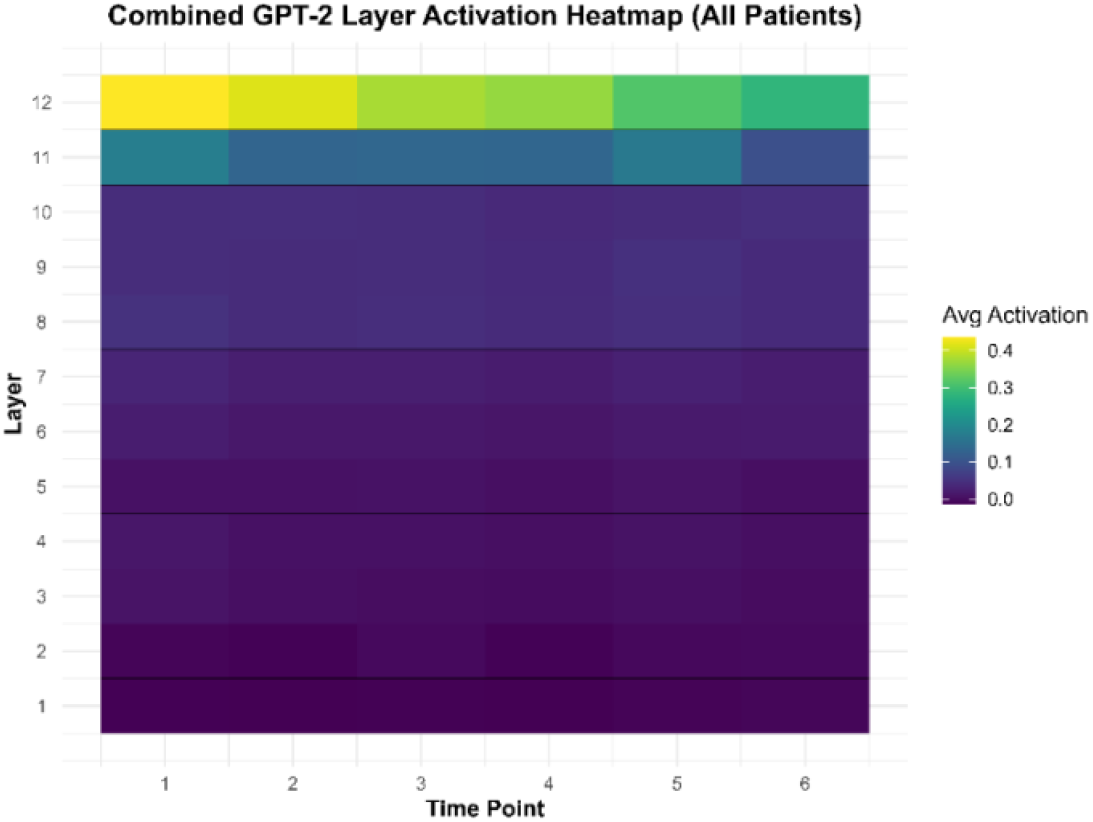
Combined GPT-2 Layer Activation Heatmap. Heatmap showing the average GPT-2 layer activation across all patients and time points. Each tile represents the mean hidden state activation for one layer (y-axis) during a specific time point (x-axis). Color intensity corresponds to activation magnitude, with deeper layers showing consistently higher activation.

### Layer Activation Patterns across Aphasia Subtypes

To assess whether GPT-2 layer wise activations varied by aphasia subtype, we conducted twelve one-way ANOVAs (one for each of the 12 layers) using aphasia subtype as the independent variable and mean activation values as the dependent variable. These analyses revealed significant main effects of subtype at all layers (all p < 0.001), indicating consistent subtype-related differences in activation magnitudes (Figure 3). No time variable was included in these models, as timepoint-averaged activations were used for this analysis. Post hoc comparisons via Tukey’s HSD test (α = 0.05) revealed specific differences between subtypes, particularly between Broca’s and Anomic aphasia and between Wernicke’s and Broca’s aphasia. A full list of comparisons and adjusted p values is provided in Appendix A Table A3.

**Figure 3.**
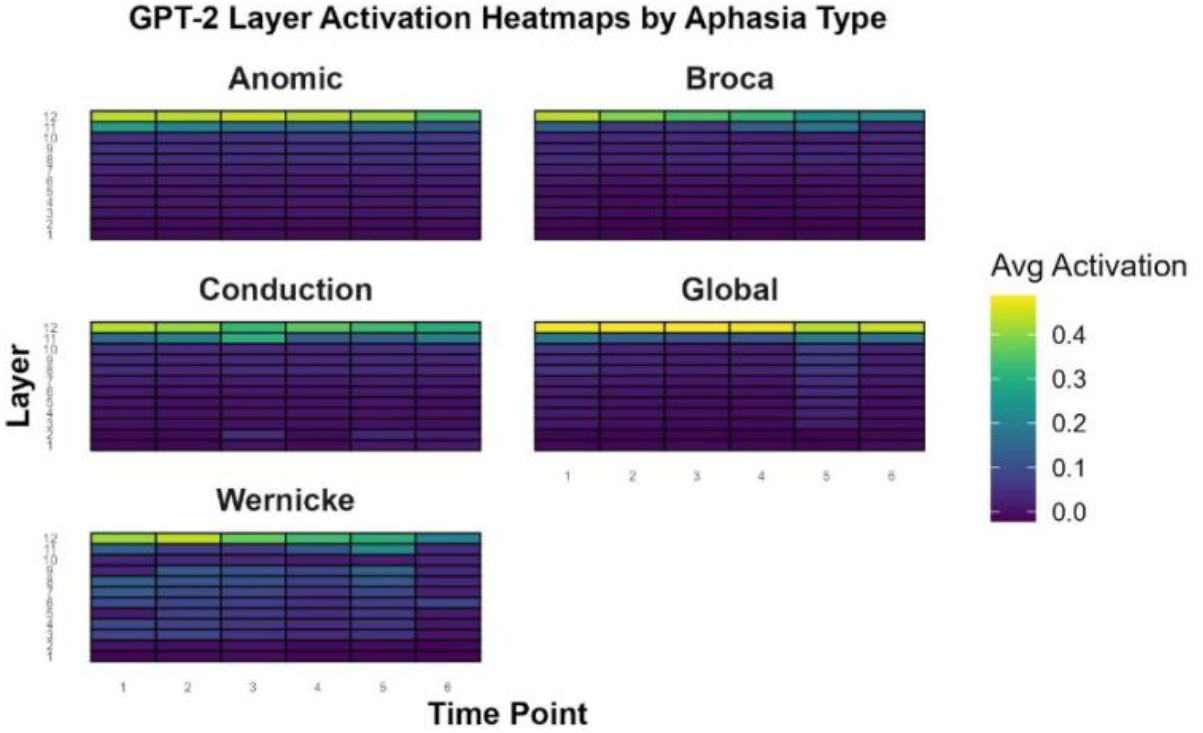
GPT-2-layer activation heatmaps by aphasia subtype. Each panel shows average activation across Time Point (x-axis) and GPT-2 layers (y-axis) for one aphasia type. Color intensity represents mean activation. Different patterns across subtypes suggest major differences in representational engagement between groups.

### Longitudinal Dynamics of GPT-2 Layer Activations During Recovery

To quantify where significant changes in layer activation occurred over time, we first fitted linear mixed-effects models for each of the 12 GPT-2 layers with time points (1-6) as fixed effects and participants as random effects. F tests on the time effect revealed that several deeper layers exhibited significant longitudinal modulation. These effects remained significant after False Discovery Rate (FDR) correction at q < .05, indicating that activation patterns in later layers changed systematically across the recovery period.

To further characterize the magnitude and direction of these effects, we conducted Wilcoxon signed-rank tests to compare activation between the first and last time points across patients, regardless of direction. The individual trajectories exhibited both increases and decreases in activation; however, the most significant group-level changes were noted in the final three layers of GPT-2. Layer 12 demonstrated the greatest effect size (r = 0.84), followed by Layer 11 (r = 0.67) and Layer 10 (r = 0.24), with all the results achieving statistical significance after adjustment for multiple comparisons. Conversely, earlier layers (Layers 1–9) presented smaller and statistically nonsignificant median effect sizes, with r values predominantly less than 0.24. Figure 4 presents the effect sizes derived from the Wilcoxon tests. The results indicated the extent of change rather than its direction, emphasizing that the deeper layers of GPT-2 experienced the most significant longitudinal modulation during the recovery period.

**Figure 4.**
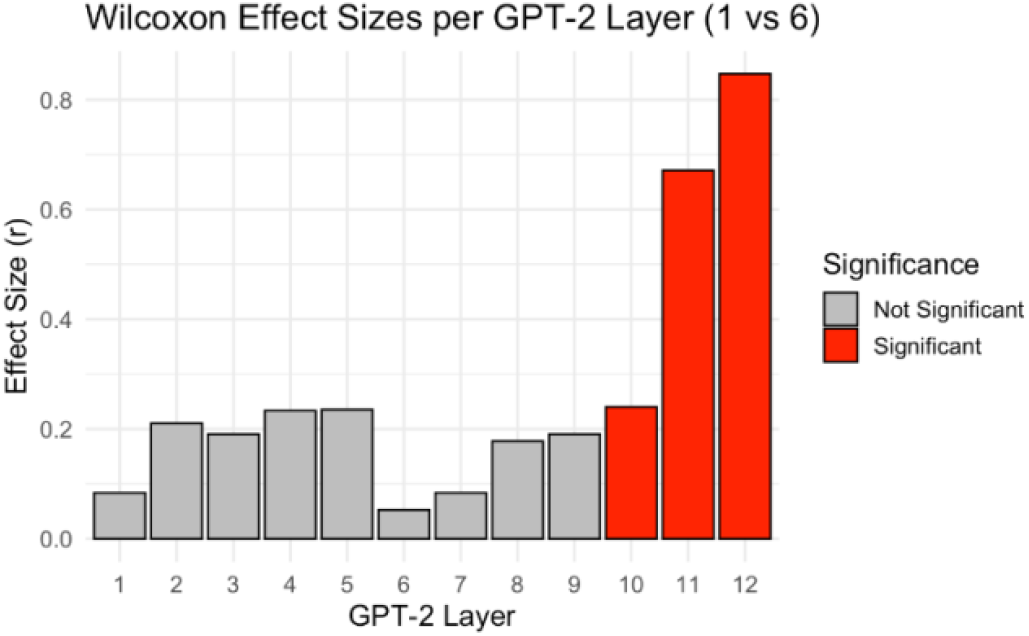
Wilcoxon Effect Sizes per GPT-2 Layer (1 vs 6) Wilcoxon effect sizes (r) for each GPT-2 layer comparing activations between Time Point 1 and Time Point 6. Bars are colored by significance after FDR correction. Deeper layers (10–12) showed the largest and statistically significant changes over time.

### Correlation with Clinical Language Scores

Layer wise correlation analyses demonstrated significant positive associations between GPT-2 activation and WAB-AQ scores across most layers, especially in the early and deep layers (e.g., Layer 2: r = 0.52, p < 0.001; Layer 10: r = 0.51, p < 0.001), suggesting that greater activation correlated with better language performance across the board. In Figure 5, layers whose activation significantly correlates with aphasia severity are marked.

**Figure 5.**
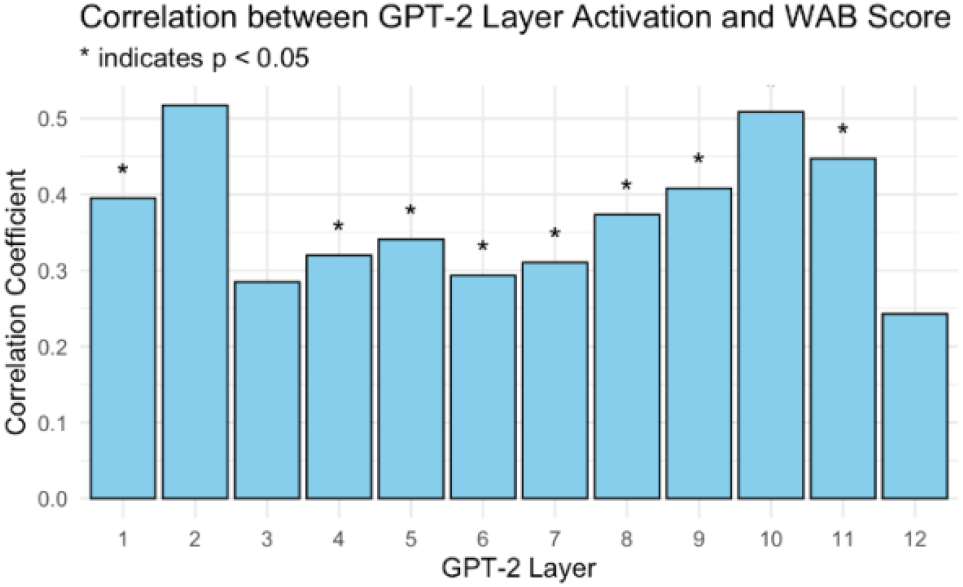
Correlation between GPT-2 Layer Activation and WAB Score. The bar plot shows the Pearson correlation coefficients between mean activation in each GPT-2 layer and patients’ WAB-AQ scores. Asterisks (*) indicate statistically significant correlations at p < 0.05. The strongest correlations were observed in the deeper layers (especially Layers 2, 10, and 11), suggesting that activation strength in deeper layers is more predictive of aphasia severity.

To better capture the relationship between aphasia severity and neural activation, we examined the continuous association between WAB-AQ scores (one per participant) and GPT-2-layer activations. Several layers, particularly the deeper ones (Layers 10–12), exhibited positive trends, suggesting a graded increase in representational strength with improved language ability. This pattern supported the idea that deeper transformer layers are more sensitive to clinical variation in aphasia severity (Figure 6).

**Figure 6.**
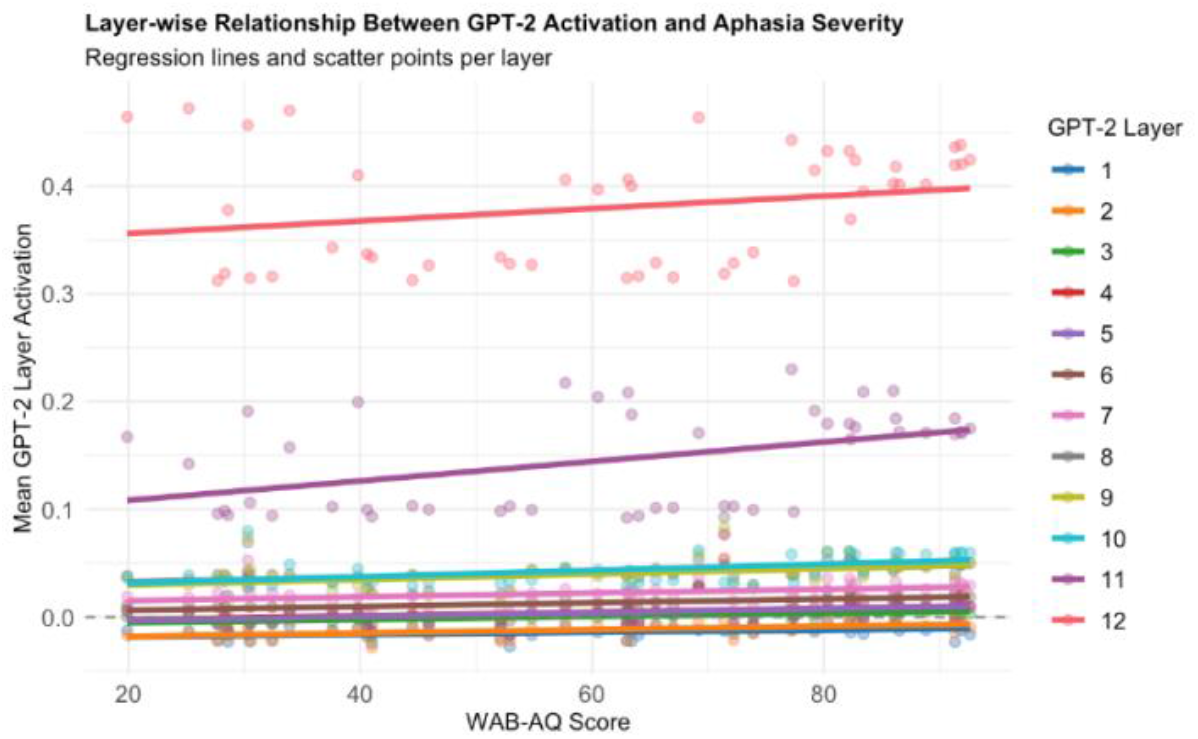
Layer wise Relationship Between GPT-2 Activation and Aphasia Severity. Each point represents an individual participant. Colored regression lines represent layerwise linear trends, with deeper layers (e.g., 10–12) exhibiting stronger positive associations between activation and aphasia severity. This visualization illustrates how GPT-2 activations scale continuously with clinical language performance without the need for categorical grouping.

## Discussion

### Overview of Key Findings

This study demonstrated that GPT-2-layer activations distinguish between aphasic and neurotypical individuals (while the broad patterns are similar), with notable differences identified in layers 1, 2, 5, 8, 10, 11, and 12. Compared with other subtypes, speakers with Broca’s aphasia presented unique activation patterns, particularly in the deeper layers. Layers 10 to 12 presented the most significant recovery-related alterations, with Layer 12 exhibiting the greatest impact size. Activation at multiple levels, specifically layers 2, 10, and 11, was positively correlated with WAB-AQ scores, indicating that enhanced model representations are associated with improved language proficiency.

### Layer Activation in Typical and Aphasic speech

Our study revealed that the general pattern of layer activation in GPT-2 during speech processing is largely consistent between aphasic and normal speakers, especially in the model’s deeper layers. However, notable differences were observed in layers 1, 2, 5, 8, 10, 11, and 12, suggesting that, compared with neurotypical controls, aphasic subjects demonstrated modified representational depth at baseline. These deviations may reflect disrupted linguistic processing or reduced semantic abstraction in aphasic speech, particularly in the deeper layers where contextual integration and higher-order meaning representations are thought to occur (Rogers et al., 2021; Tenney et al., 2019). Early layer differences (layers 1 and 2) may also suggest abnormalities in token-level or syntactic processing, which is consistent with prior evidence that aphasia affects both lower- and higher-order stages of language representation within the model (Jawahar et al., 2019).

In narrative-discourse samples of people with aphasia, the changes in hidden-state activation in deeper layers may reflect how the language model attempts to preserve semantic coherence in response to structurally impaired input (Caucheteux et al., 2021; Schrimpf et al., 2021). Despite the superficial inconsistencies in aphasic speech, the deeper layers of the GPT-2 continue to engage their semantic representation mechanisms, indicating that the model still attempts to construct discourse-level meaning from impaired input. Importantly, activation at these layers does not necessarily imply a correct or successful interpretation; rather, it reflects that the model continues to process and integrate available linguistic cues at the semantic level.

### Subtype-Specific Patterns

Layer wise analyses revealed that most significant longitudinal changes occurred in the deepest layers of GPT-2, particularly Layer 12. These layers showed consistent decreases in activation over time at the group level (except in Wernicke’s aphasia), despite individual variability. As these layers are hypothesized to integrate abstract semantic and contextual information, their decreased engagement may reflect growing discourse cohesion or reduced reliance on compensatory effort as patients recover.

Importantly, however, changes in layer activation are open to multiple interpretations. Higher activation could indicate that the system is working harder at a given level of analysis to manage impaired input, whereas lower activation may suggest reduced processing demand or greater efficiency. Conversely, increased activation might also reflect that the system is successfully extracting more useful information from the input. Thus, activation changes should be interpreted cautiously, as they may signal either compensatory effort or improved processing efficiency depending on context.

While all subtypes demonstrate a general trend toward greater activation in deeper GPT-2 layers, which is consistent with their role in semantic integration and discourse-level processing (Caucheteux et al., 2021; Tenney et al., 2019), pairwise statistical comparisons revealed distinct subtype-specific activation profiles across these layers.

The differences in layer-activation levels between different aphasic subtypes (by WAB-R classification) allow us to speculate on the possible nature of these patterns. Notably, as detailed in the results section, individuals with Broca’s aphasia exhibited significantly greater activation in the deeper layers of GPT-2 (10–12) than other subtypes did. This persistent activation pattern may reflect the model’s continued sensitivity to semantic-level cues even in syntactically sparse input. These changes and persistent deep-layer engagement may reflect the model’s continuous attempt to semantically reconstruct meaning from syntactically sparse and effortful speech. The stability of this response suggests that, despite grammatical disruptions, semantic content remains sufficiently recoverable to engage GPT-2’s deeper semantic layers.

In contrast, Wernicke’s aphasia exhibited markedly different activation profiles, with significantly lower deep-layer activation than Broca’s aphasia and a more even distribution across the middle to deep layers. This dispersed pattern may correspond to the fluent but semantically incoherent nature of Wernicke’s speech, which engages GPT-2’s higher-order representations inconsistently. The relative flatness in activation over time further supports limited structural compensation or adaptive reorganization in this group.

Anomic and conduction aphasia both showed a gradual decrease in activation in Layers 11–12, but conduction aphasia consistently exhibited significantly greater activation than did anomic aphasia across several layers. This may reflect a greater need for phonological monitoring and semantic reinterpretation in conduction aphasia, as patients attempt to self-correct disrupted output. This evolving pattern may reflect ongoing recovery or refinement in discourse formulation as lexical retrieval and error correction improve (Halai et al., 2017).

Global aphasia exhibited early deep-layer activation, potentially indicating GPT-2’s attempt to derive meaning from significantly diminished or impoverished input. Nonetheless, this activation exhibited minimal variation and only a slight decrease over time, indicating stabilization or limited semantic recovery at a basic communicative level.

The subtype-specific patterns indicate that GPT-2 captures broad trends in semantic integration across aphasia while also identifying nuanced differences in the engagement of subtypes with deep layers over time. This finding indicates that the model’s capability is an effective instrument for analyzing discourse-level disruptions in aphasic speech.

In addition, our analysis indicates that the internal activations of the GPT-2 vary among aphasia subtypes and systematically correspond to individual language proficiency, as measured by WAB-AQ scores. The observed positive correlations in both the surface and deeper layers indicate that surface-level and integrative semantic processes encoded in GPT-2 play a role in capturing linguistic impairment. The group-level comparisons support this assertion, demonstrating enhanced activations in patients with elevated WAB scores, especially in deeper layers, aligning with prior assertions that later transformer layers represent higher-order linguistic structures.

These findings may serve as a clinical reference, endorsing the application of transformer-derived activations as a complementary or even alternative indicator for language severity and emphasizing their potential in low-burden diagnostic tools.

### Limitations and Future Directions

A major limitation of the present study is that although GPT-2 has demonstrated alignment with brain data, the precise linguistic functions of its individual layers remain somewhat speculative. Our analyses depend on the current literature connecting deeper layers to discourse-level semantics; nonetheless, further controlled investigation is needed. The speech transcripts did not include acoustic or prosodic data, which could be valuable in future multimodal methodologies. Future research should incorporate neurotypical comparison groups, broaden analysis to multilingual environments, and utilize layer probing approaches to more accurately correlate linguistic deficiencies with model internals. Future research should prioritize and integrate automated speech recognition techniques with large language models.

## Conclusion

This research presents a new approach for the automated and objective analysis of aphasic speech via transformer-based language model representations. This method circumvents the need for manual coding by utilizing only textual transcripts, providing extensive analysis of discourse-level patterns in aphasia. This study suggests that the internal activations of GPT-2 can reflect subtype-specific linguistic profiles, monitor changes during recovery, and categorize patients without requiring external labels. These findings provide an early indication of the viability of employing model-derived metrics as biomarkers for diagnosis and monitoring progress in clinical environments. Additional research is necessary to elucidate the distinct functional roles of each model layer, which may facilitate targeted computational evaluations and tailored language interventions.

## Data Availability Statement

The dataset analyzed in this study consists of identifiable clinical speech recordings collected from individuals with post-stroke aphasia. Due to the presence of potentially identifiable personal and health-related information, the raw speech data cannot be made publicly available in order to protect participant privacy and confidentiality. Processed numerical activation matrices derived from the speech data, along with all analysis scripts used in this study, are available from the corresponding author (Seyed Saeed Ahmadi) upon reasonable request, subject to approval by the relevant ethics committee.

## Notes

### Competing Interest Statement

The authors have declared no competing interest.

